# Conserving evolutionary history to safeguard our future: incorporating the Tree of Life into biodiversity policy

**DOI:** 10.1101/2021.03.03.433783

**Authors:** Rikki Gumbs, Abhishek Chaudhary, Barnabas H. Daru, Daniel P. Faith, Félix Forest, Claudia L. Gray, Aida Kowalska, Who-Seung Lee, Roseli Pellens, Sebastian Pipins, Laura J. Pollock, James Rosindell, Rosa A. Scherson, Nisha R. Owen

**Affiliations:** EDGE of Existence programme, Zoological Society of London, London, NW1 4RY, United Kingdom; Imperial College London, Silwood Park Campus, Buckhurst Road, Ascot, Berkshire, SL5 7PY, United Kingdom; Department of Civil Engineering, Indian Institute of Technology (IIT) Kanpur, 208016 Kanpur, India; Department of Life Sciences, Texas A&M University-Corpus Christi, Corpus Christi, Texas, 78412, USA; School of Philosophical and Historical Inquiry, The University of Sydney, Sydney, Australia; Royal Botanic Gardens, Kew, Richmond, Surrey, TW9 3AE, United Kingdom; On the EDGE Conservation, 152a Walton Street, Chelsea, SW3 2JJ; Environmental Assessment Group, Korea Environment Institute, Sejong, 30147, Republic of Korea; Institut de Systématique, Evolution, et Biodiversité (Muséum national d’Histoire naturelle, Centre National pour la Recherche Scientifique, Sorbonne Université, Ecole Pratique de Hautes Etudes, Université des Antilles) CP 50, 45 rue Buffon - 75005 Paris, France; Department of Biology. McGill University. Montréal, Québec, H3A 1B1, Canada; Departamento de Silvicultura y Conservación de la Naturaleza, Universidad de Chile., PO Box 9206, Santiago, Chile

## Abstract

Following our failure to fully achieve any of the 20 Aichi biodiversity targets, the future of biodiversity rests in the balance. The Convention on Biological Diversity’s Post-2020 Global Biodiversity Framework (GBF) presents us with the opportunity to preserve Nature’s Contributions to People (NCPs) for current and future generations through conserving biodiversity and averting extinction across the Tree of Life. Here we call attention to our need to conserve the Tree of Life to maintain its benefits into the future as a key mechanism for achieving intergenerational equity. We highlight two indicators available for adoption in the post-2020 GBF to monitor our progress towards safeguarding the Tree of Life. The Phylogenetic Diversity indicator, adopted by IPBES, can be used to monitor biodiversity’s capacity to maintain NCPs for future generations. The EDGE (Evolutionarily Distinct and Globally Endangered) Index monitors how well we are performing at averting the greatest losses across the Tree of Life by conserving the most distinctive species. By committing to safeguarding the Tree of Life post-2020, we can reduce biodiversity loss to preserve nature’s contributions to humanity now and into the future.

## Biodiversity policy and the Tree of Life

Existing policy has failed to stem biodiversity declines across the board (Díaz et al. 2019), partially achieving only six of the 20 Aichi biodiversity targets (Secretariat of the Convention on Biological Diversity 2020a). As a result, it is clear that only by being highly ambitious can we have any chance of improving the outlook for global biodiversity by 2050 (Díaz et al. 2020). While parties of the Convention on Biological Diversity (CBD) work towards the post-2020 Global Biodiversity Framework (GBF), it is vital that goals and targets for the coming decades are not only ambitious in the amount of biodiversity (species, areas, ecosystems and their contributions to people) to be protected, but also that they consider the many biodiversity facets and the interlinks among them.

A critical and often overlooked facet of biodiversity is the evolutionary history represented by a set of species across the Tree of Life, measured by Phylogenetic Diversity (PD; Faith 1992). PD approximates the entire suite of features shared by, and unique to, the species in a given set by measuring the branches of the Tree of Life that connect them. Consequently, the greater the loss of PD, and therefore of evolutionary history, the more varied and distinct features we stand to lose (Faith 1992). As a measure of living variety, PD captures both the current benefits and as yet unexplored options for humanity provided by biodiversity, or the option values of biodiversity (Faith 2016).

Empirical studies have shown that by focusing on PD we protect a larger diversity of benefits from biodiversity than with random or taxonomic based choices (Forest et al. 2007; IPBES 2019; Molina-Venegas 2021; Molina-Venegas et al. 2021). For example, Forest et al. (2007) illustrated that maximising the distribution across the Tree of Life when selecting sets of plant species in the Cape of South Africa would more efficiently retain plants with known benefits than selecting based on generic richness alone. Similarly, Molina-Venegas et al. (2021) showed that maximising the branches of the Tree of Life more efficiently safeguarded a larger number and wider variety of benefits to people (hereafter ‘benefits’)—including materials, food, medicine and fuels—provided by the world’s plants than with a random selection.

Maintaining the benefits and options generated by biodiversity for future generations (the biodiversity option value represented by PD; Faith 2018), as a key mechanism for achieving intergenerational equity, is particularly important in the context of a changing environment and the challenges that biodiversity faces going forward (Pollock et al. 2017; IPBES 2019; Veron et al. 2019; Molina-Venegas et al. 2021; Robuchon et al. 2021). Indeed, Díaz et al. (2020) recognise that, if global policy is to ‘bend the curve’ of biodiversity loss whilst securing a broad range of benefits, we must set and attain highly ambitious goals that include prioritising the conservation of evolutionarily distinct lineages to effectively safeguard the Tree of Life.

Accordingly, there is increasing focus on incorporating PD into national (e.g. Creswell & Murphy 2016) and global biodiversity policy initiatives (UNEP-WCMC 2015; IPBES 2019; Secretariat of the Convention on Biological Diversity 2021). The Intergovernmental Science-Policy Platform on Biodiversity and Ecosystem Services (IPBES) adopted the status of threatened PD as an indicator (hereafter IPBES PD indicator) for Nature’s Contributions to People (NCPs; Díaz et al. 2019; IPBES 2019): linking PD to the maintenance of options (the overall capacity of biodiversity to support a good quality of life into the future; NCP 18), and thus the continued provision of medicinal, biochemical and genetic resources (NCP 14), and learning and inspiration (NCP 15; Díaz et al. 2019; IPBES 2019). Initial estimates of the IPBES PD indicator have been produced for several clades (amphibians, corals, cycads, birds, mammals and squamates) as part of IPBES’s regional and global assessments (Faith et al. 2018; IPBES 2018; Martín-López et al. 2018).

For global biodiversity policy to effectively protect the Tree of Life and the benefits it provides, we need a particular focus on distinctive species that embody a disproportionate amount of threatened evolutionary history. The extinction of highly evolutionarily distinctive species, with few close relatives on the Tree of Life, results in the irreversible loss of not only their unique characteristics but also their potential benefits, and should be avoided (Rounsevell et al. 2020; Díaz et al. 2020). The IUCN has long recognised the importance of conserving distinctive species (IUCN 1980) and resolved in 2012 to halt the loss of evolutionarily distinct species (IUCN 2012). Evolutionarily distinct species are increasingly recognised as of particular conservation importance globally (UNEP-WCMC et al. 2018; Carta et al. 2019; Díaz et al. 2020), and their protection will play an important role in safeguarding future benefits derived from biodiversity (Molina-Venegas 2021).

Priority Evolutionarily Distinct and Globally Endangered (EDGE) species are those species that are threatened with extinction and represent large contributions to the PD expected to be lost from a taxonomic group (above median; Isaac et al. 2007), and should therefore be prioritised for conservation action. The EDGE approach has been widely applied to various groups of animals (Isaac et al. 2007, 2012; Jetz et al. 2014; Curnick et al. 2015; Owen et al. 2015; Stein et al. 2018; Gumbs et al. 2018) and plants (Daru & le Roux 2016; Forest et al. 2018), leading to the identification of sets of species that are both highly irreplaceable and vulnerable. EDGE species are the focus of the Zoological Society of London’s EDGE of Existence programme, which has funded more than 120 conservation projects worldwide and maintains publicly available priority lists for numerous taxonomic groups (ZSL EDGE of Existence 2022).

Here, we provide expanded details on two indicators designed to monitor the conservation status of PD and the most distinctive species across the Tree of Life, and apply them to the world’s birds, mammals, and cycads. First, we outline an updated approach to the IPBES PD indicator to monitor overall trends in the expected loss of PD. Second, we outline the EDGE Index, a species-focused indicator that tracks the change in conservation status for the most evolutionarily distinctive and threatened species through time, to highlight species whose conservation can safeguard large amounts of threatened evolutionary history. We discuss how these indicators complement—and expand on—existing tools for monitoring biodiversity, and how they can be incorporated into existing policy frameworks—such as the CBD—at national and global scales (see Appendix S1).

## PD indicator

We propose an update to the existing IPBES PD indicator, improving its robustness and applicability for biodiversity policy at regional and global scales, including in the post-2020 GBF. Specifically, this update incorporates standardised IUCN Red List extinction risk data for all species in taxonomic groups where available, for multiple time points where applicable (IUCN 2020; Henriques et al. 2020), to generate trends in the status of PD using the established expected PD loss framework (Faith 2008; Faith et al. 2018).

Expected loss is an established probabilistic approach to measuring the status of diversity (Witting & Loeschcke 1995; Weitzman 1998) that is now widely applied to inform both biodiversity monitoring, including the IPBES PD indicator (Faith et al. 2018; Davis et al. 2018; IPBES 2019; Gumbs et al. 2020) and conservation action (Redding & Mooers 2006; Isaac et al. 2007; Steel et al. 2007; Nunes et al. 2015; Forest et al. 2018). The expected loss of PD is the amount of evolutionary history expected to be lost in a given amount of time based on the current extinction risks faced by the set of species (Faith 2008).

When an ancestral branch of the Tree of Life has several descendant species that are all at low risk of extinction, the risk of losing that branch is relatively low, because it would require all descendants to become extinct. However, as the probability of extinction increases for each descendant species, the associated risk of losing the ancestral branch that they share also increases. This interaction between related species’ risk of extinction is known as ‘PD complementarity’ (Faith et al. 2004). The greater the proportion of long branches of the Tree of Life that have only threatened species as descendants, the greater the expected loss of PD. For example, the branch that connects pangolins to all other mammals is long and continues to exist only while the eight descendent species of pangolin survive. At present all eight are threatened, together representing an entire mammalian order at risk of extinction.

Our updated approach to estimate the PD indicator explicitly incorporates the concept of PD complementarity to directly estimate the expected loss of branches of the Tree of Life. This approach is widely recommended for accurate calculation of expected PD loss (Steel et al. 2007; Faith 2008; Nunes et al. 2015; Faith et al. 2018; Gumbs et al. 2022) but was not used for the initial IPBES PD indicator formulation (for detailed methods, see Appendix S2). Our updated PD indicator utilises probabilities of extinction—converted from IUCN Red List categories (Mooers et al. 2008)—to calculate the amount of PD expected to be lost from each branch of the Tree of Life, based on the probability that the set of species responsible for the survival of that branch are lost in the future. We assign increasing values of extinction risk to Red List categories of increasing severity, rather than treating all threatened species at equal risk of extinction (as in the IPBES ED approximation of Faith et al. 2018). Thus, our updated PD indicator is better aligned with the usage of extinction risk data in the Red List Index (RLI; Butchart et al. 2004), another key biodiversity indicator.

We demonstrate the utility of this updated PD indicator using three taxonomic groups: birds, mammals and cycads. Given increased levels of extinction risk through time for each of the three taxonomic groups, trends in their expected PD loss are worsening (Figure 1). Though the magnitude of expected PD loss varies across extinction risk quantifications, the trends in relative change through time are consistent (within 10% across all groups; see Appendix S2).

**Figure 1:**
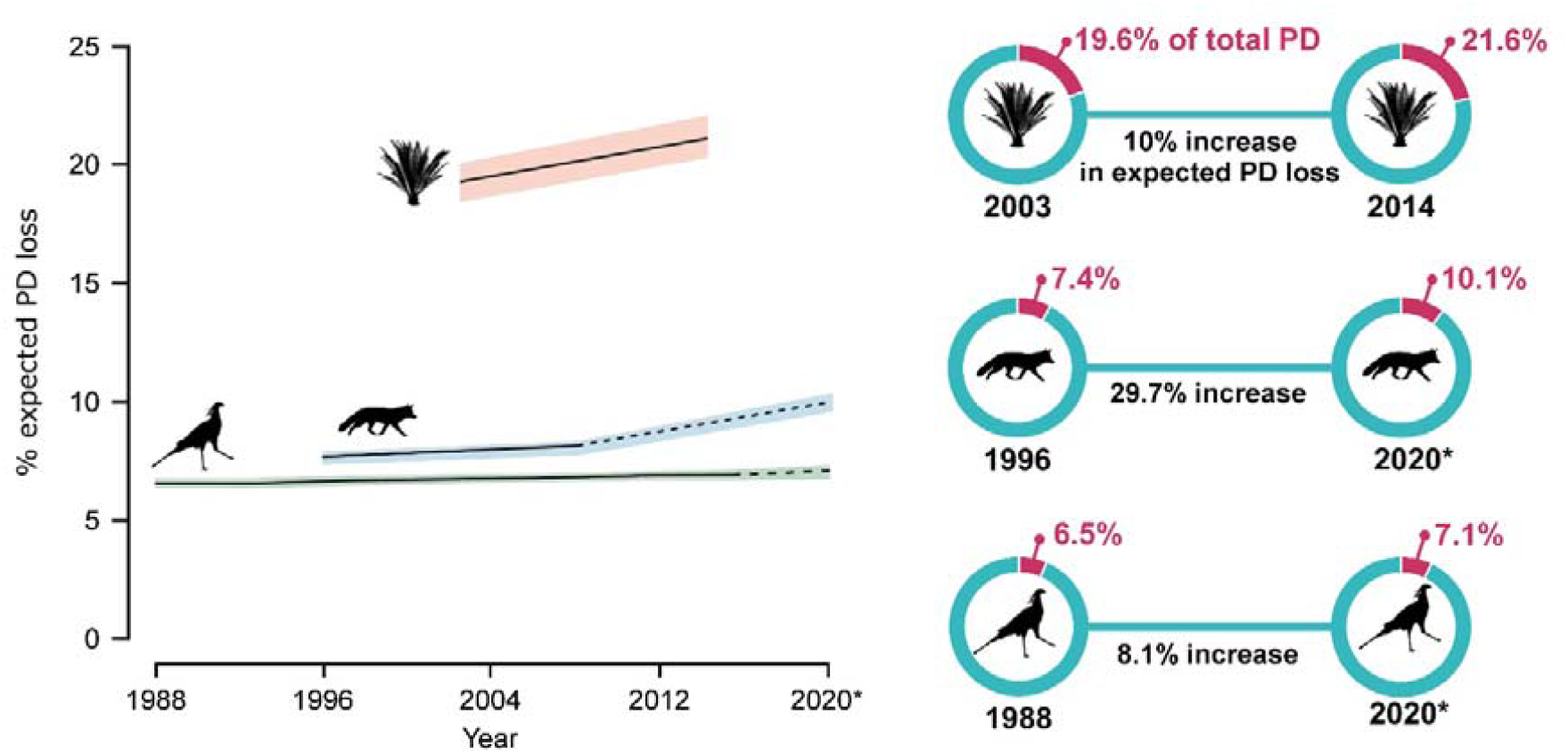
The PD indicator: tracking PD loss through time. Left panel: trends in percentage of expected PD loss for the world’s mammals (blue), birds (green) and cycads (pink), based on current and historical IUCN Red List assessments. Right panel: detail of this change, where each circle represents the total PD of the clade, and the percent expected loss is coloured in red, for the baseline (left circle) and latest (right circle) time points, with the percent change of expected PD loss between each time point given under the connecting line. *The 2020 timepoints displayed are not official Red List Index (RLI) timepoints of comprehensive assessments for all mammals and birds—as these are not yet available–but represent the latest status of these assessments for both clades (see Appendix S2); the trendlines from official RLI data to these 2020 timepoints are therefore dashed. The shaded regions around each trend line represent the range of values. There are insufficient repeated Red List assessments to produce a 2020 timepoint for cycads.

We also present the baseline PD indicator values for these groups calculated using the ‘EDGE’ extinction risk quantification (where probability of extinction halves with every decrease in Red List category, from 0.97 for Critically Endangered to 0.485 for Endangered and so forth, following Mooers et al. (2008), see Appendix S2 for detailed methods). This calculation aligns with the calculation of EDGE scores for identifying priority species (see ‘EDGE Index’ section, below). Our approach provides the most conservative estimates of relative change for each group (see Appendix S2 for baselines using other extinction risk quantifications). Mammals have the greatest projected relative increase in expected PD loss (median = 29.7% increase from 1996-2020) though, due to their elevated extinction risk, cycads (21.6%) are currently expected to lose more than double the proportion of PD than mammals (10.1%), and three times that of birds (7.1%; Figure 1).

By comparison, the initial IPBES PD indicator was derived from the Evolutionary Distinctiveness scores (ED; which shares the PD of each branch of the Tree of Life equally amongst all descendants; Isaac et al. 2007) of threatened species (Vulnerable, Endangered and Critically Endangered on the IUCN Red List), to estimate the proportion of imperilled PD (Faith et al. 2018). The initial IPBES PD indicator estimated levels of expected PD loss to around 2x higher than those from the most pessimistic probabilities of extinction applied here (Appendix S2). This is to be expected given the initial IPBES PD indicator did not incorporate PD complementarity, which has now been included in our advanced version to more accurately estimate the expected loss associated with branches with multiple descendants.

Considering Phylogenetic Diversity’s link to the provision of current and future benefits (Forest et al. 2007; IPBES 2019; Molina-Venegas et al. 2021), and its adoption by IPBES to indicate the capacity of biodiversity to keep options open in order to support a good quality of life (Faith et al. 2018; Díaz et al. 2019), it provides a unique and versatile tool with which to monitor Nature’s Contributions to People while maintaining intergenerational equity. The PD indicator is therefore relevant to all aspects of Goal B of the draft Global Biodiversity Framework that relate benefits for people to biodiversity (Secretariat of the Convention on Biological Diversity 2020b, 2021).

## EDGE Index

An established tool for identifying evolutionarily distinct species whose conservation should be prioritised to avoid particularly large losses of PD is the EDGE approach (Isaac et al. 2007). The EDGE Index utilises the EDGE framework to monitor the conservation status of—and the impact of conservation action for—highly evolutionarily distinct and threatened species through time. The EDGE Index deconstructs existing RLI extinction risk data for the world’s most evolutionarily distinct and threatened species to provide explicit monitoring of conservation successes and failures, extinctions, and changes in expected PD loss for these irreplaceable sets of species.

The EDGE scores of species can be calculated as species-specific contributions to expected PD loss (Steel et al. 2007; Faith et al. 2018; Gumbs et al. 2022), and can be derived for birds, cycads and mammals from the phylogenetic trees with branches weighted by extinction risk for the calculation of the expected PD loss indicator outlined above. The expected PD loss contribution of a species can be calculated from a phylogenetic tree whose branch lengths have been multiplied by the overall probability of extinction of descendant species, such as those trees used to calculate the PD indicator outlined above. For each species the lengths of the phylogenetic branches connecting the species at the tip to the root of the tree are summed. Although Díaz et al. (2020) proposed a novel measure of evolutionary distinctiveness by which to prioritise PD, this approach does not perform as well as existing distinctiveness measures—such as that underpinning the EDGE approach— at capturing PD (Appendix S2).

In order to calculate our example EDGE index values, we first calculated EDGE scores for birds, cycads and mammals from the 100 phylogenetic trees—with branches weighted by extinction risk. We used the median EDGE score for each species to represent its EDGE score at that point in time. Priority EDGE Species were identified as those species with above median expected PD contributions and in Red List threatened categories at that RLI time point (Isaac et al. 2007; Gumbs et al. 2022). The whole process was repeated for each RLI time point and we calculated several components as follows (also see Appendix S2):

a. i. Changes in the number of EDGE species, increasing as more highly distinctive species become threatened, or decreasing as highly distinctive species move into non-threatened Red List categories;

a. ii. Changes in the amount of associated expected PD loss according to changes in EDGE species extinction risk, indicating the effectiveness of conservation efforts in averting the greatest losses of PD;

a. iii. Number of EDGE species, from a previous timepoint, that have since been declared in an ‘extinct’ Red List category: Critically Endangered (Possibly Extinct); Extinct in the Wild; or Extinct;

b. Changes in the extinction risk of EDGE species, those moving into worse Red List categories (‘uplistings’) indicate insufficient conservation efforts for the most distinctive and threatened species, whereas increased numbers of EDGE species moving into less severe Red List categories (‘downlistings’) indicate effective conservation efforts for these species.

The number of EDGE species increased for all three taxonomic groups between their first and most recent Red List Index assessments (Figure 2). Further, the conservation status of EDGE species is deteriorating, with greater numbers of uplistings than downlistings for each group, most notably for EDGE mammals, where 56 EDGE species (12%) increased in extinction risk between 1996 and 2008 (Figure 2b), compared with just 7% of all threatened mammals. Finally, each group has had at least one species transition to an ‘extinct’ category, including the Baiji (*Lipotes vexillifer*) and Spix’s Macaw (*Cyanopsitta spixii*).

**Figure 2:**
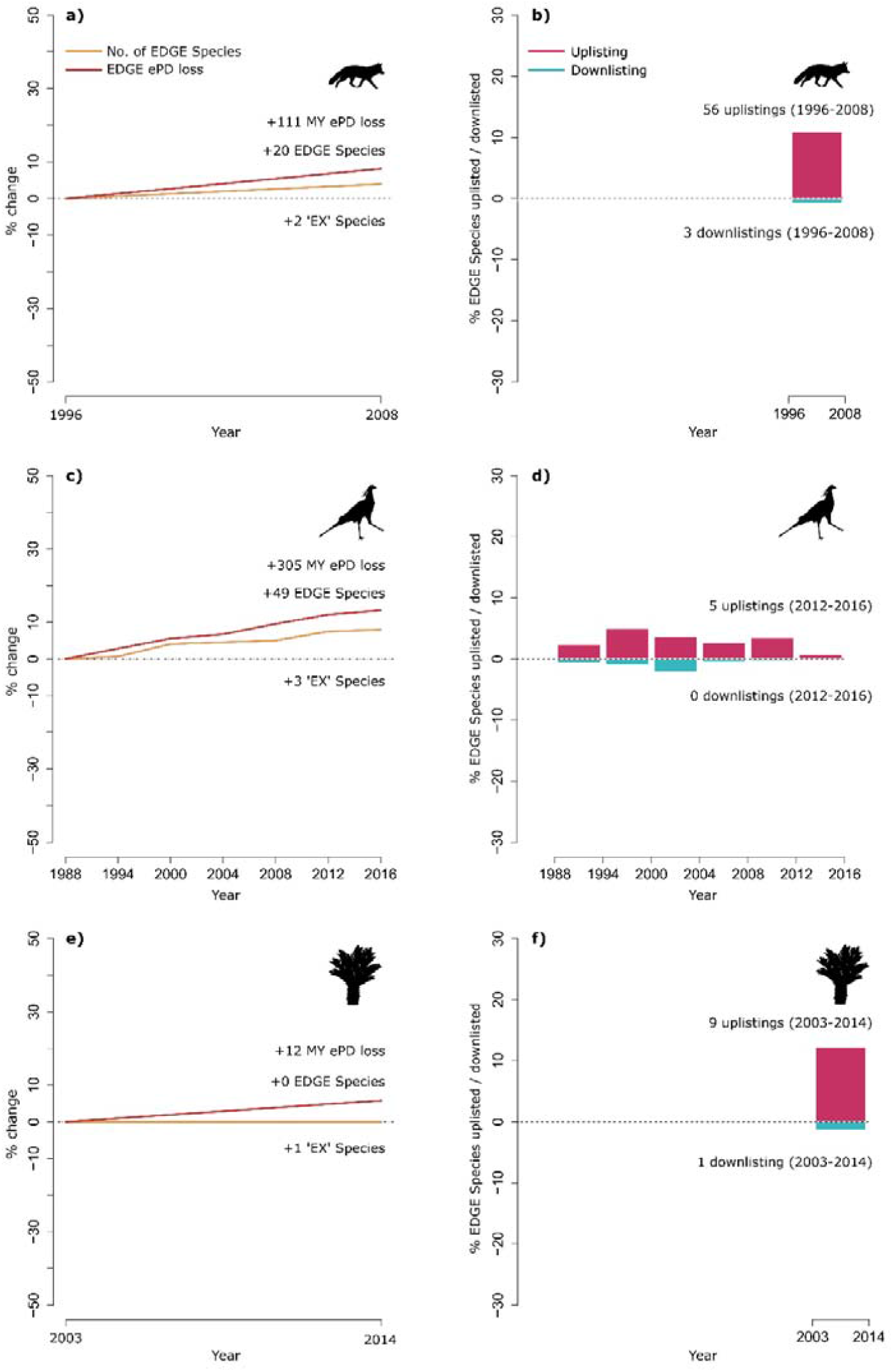
The EDGE Index: monitoring trends in extinction risk for priority EDGE Species. Left panels: tracking changes through time in the total number of EDGE species, associated expected PD loss (ePD loss), and extinctions (EX Species), of priority EDGE Species per clade; and (right panels) the changes in extinction risk (uplistings and downlistings: species moving into higher or lower Red List categories) within sets of EDGE Species, for: a-b) mammals, c-d) birds, and e-f) cycads. Changes in total number of EDGE species, associated expected PD loss, and extinct species, are cumulative from baseline timepoint (dotted line). Number of uplistings and downlistings is for each time period between time points.

This indicator complements existing broader species measures, meeting the need to prioritise evolutionarily distinct species to conserve the Tree of Life as part of any efforts to reduce extinction rate and risk (Secretariat of the Convention on Biological Diversity 2020b, 2021), making it relevant to the preventing extinctions component of draft Goal A and any proposed improvements (Williams et al. 2020).

## Conserving the Tree of Life post-2020

### Tree of Life and the Post-2020 GBF

Despite the increased recognition of PD as an integral component of biodiversity (Soto-Navarro et al. 2021), and growing evidence for its direct link to the provision of benefits (Forest et al. 2007; Molina-Venegas et al. 2021), explicit reference to PD—and the maintenance of future options—has to date been largely missing from biodiversity policy. Conservation discourses have traditionally focused on ecosystems, species, and populations—overlooking PD—but this risks isolating the conservation of biodiversity from recognition of the benefits and option values that it provides, by simply assuming these values will be maintained under current conservation paradigms. For example, without considering PD it is possible to select sets of threatened species for conservation that would actually fail to avert the greatest losses across the Tree of Life, thus reducing the overall capacity for biodiversity to provide future benefits (Figure 3; see ‘Not all extinctions are equal’ in Appendix S2).

**Figure 3:**
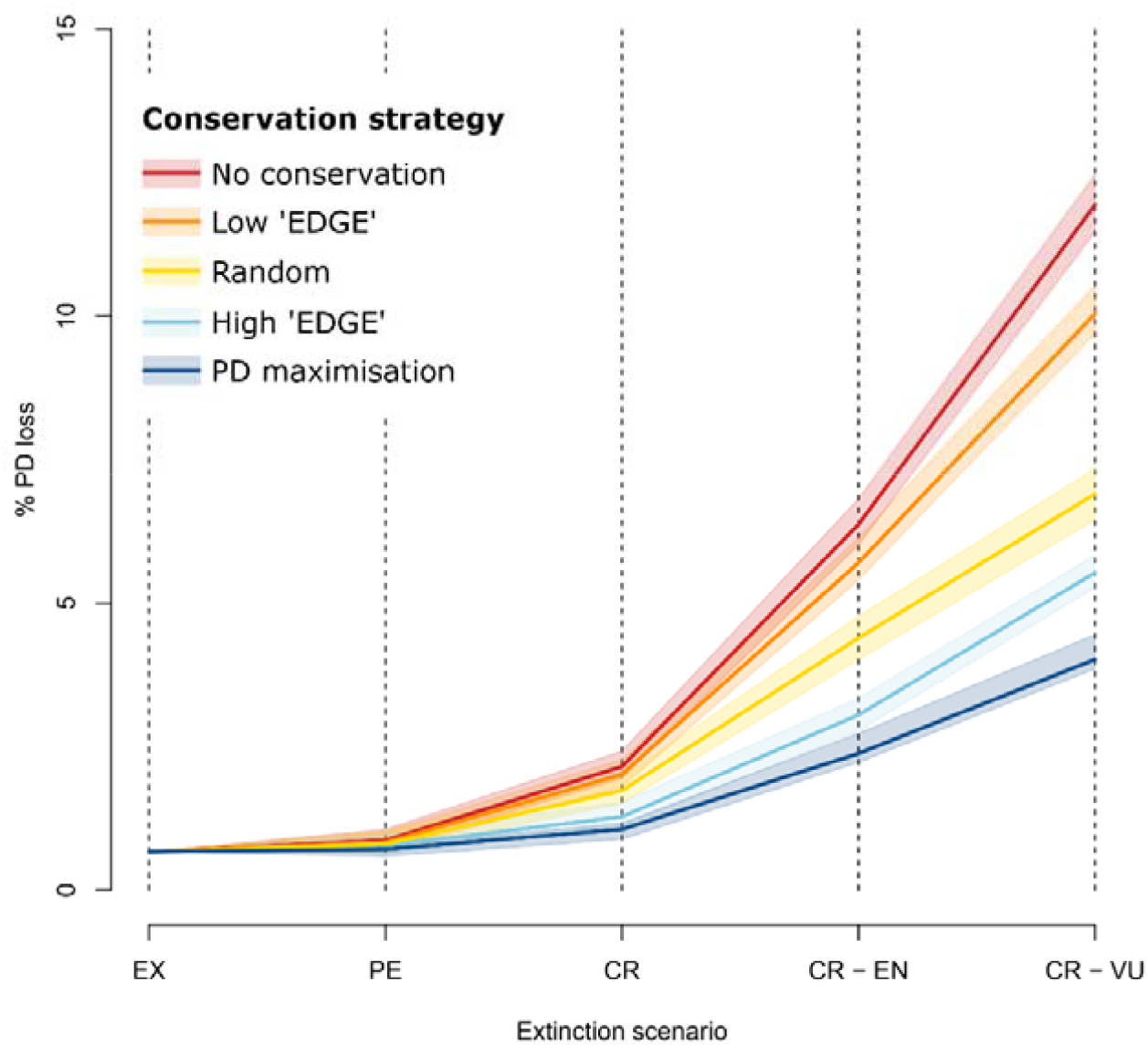
Variation in magnitude of expected loss of Phylogenetic Diversity (PD) under different conservation strategies selecting subsets of species, across four extinction scenarios for the world’s mammals: ‘PE’: only Possibly Extinct and Extinct species on the IUCN Red List are lost; ‘CR’: Critically Endangered species are also lost; ‘CR+EN’: all Endangered species are also lost; ‘CR+EN+VU: all Vulnerable species are also lost. Coloured lines represent median values of PD lost under each of the five conservation strategies across each extinction scenario. “No conservation” = no species are conserved under each extinction scenario; “low ‘EDGE’” = species in lowest quartile of expected PD loss contributions are conserved [low-ranking ‘EDGE’ Species] as a non-PD informed conservation model; “random” = a random set of species from those that meet the extinction scenario criteria, equal in size to one quartile, is conserved as our null model; “high ‘EDGE’” = species in uppermost quartile of expected PD loss contributions are conserved [high-ranking ‘EDGE’ Species] as a PD-informed conservation model; “PD maximisation” = the ideal selection of species that optimise PD is conserved. – based on a greedy algorithm. PD loss was calculated across 100 mammalian trees–- see Appendix S2 for detailed methods.

A notable exception is the acknowledgement by IPBES of option value as an important value provided by biodiversity (Díaz et al. 2015), and the subsequent explicit adoption of PD as a direct indicator to capture this and other values (IPBES 2019). This embrace of PD and option value, and the ability to report the PD indicator at global and disaggregated regional levels (IPBES 2018), provides a solid foundation for the wider adoption of PD into biodiversity policy.

While PD can evidently provide a useful tool for the monitoring of biodiversity and its associated values for a myriad of global, regional and national policy frameworks, the most pertinent and overarching of these is the CBD’s post-2020 Global Biodiversity Framework (GBF). Within the GBF, our two proposed indicators are essential for filling existing gaps (see Appendix S1). The currently listed headline indicator for Goal B (National Environmental Economic Accounts of Ecosystem Services) relates only to physical and monetary ecosystem services and assets, neglecting an entire set of non-monetary benefits and options that biodiversity provides, which must be secured (Secretariat of the Convention on Biological Diversity 2020b). While intergenerational equity is an important ambition in the CBD’s post-2020 GBF, there is no other indicator aside from PD which explicitly links the maintenance of biodiversity to the provision of benefits for future generations in a changing world. Further, none of the multiple species-focused indicators within Goal A account for between-species diversity and the features and traits that arise as a result of our planet’s evolutionary history, instead capturing only information per species (e.g. Red List Index and Genetic Diversity indicators; Secretariat of the Convention on Biological Diversity 2020b).

### National disaggregation

Given the need for nations to quantify their own values and monitor their individual progress towards both national and global targets (Rounsevell et al. 2020), it is essential that biodiversity indicators can be disaggregated to regional and national levels. In fact, the capability to monitor the status of global values of biodiversity at a national level is particularly important given that current local-scale conservation efforts can often neglect these species (Owen et al. 2019). Indeed, the benefits to humanity bestowed by species of medicinal importance, or those species that inspire awe for the natural world, transcend both political borders and generations. National responsibility for the preservation of biodiversity is particularly vital given the high numbers of species that are endemic to individual countries (Pitman & Jørgensen 2002).

Both the indicators proposed here can be effectively disaggregated to national levels. To illustrate the simplicity of this, we generated the expected PD loss indicator and EDGE Index (all components) for the birds of Kenya (Figure 4, see Appendix 2), just one example of a biodiverse country. We calculate the national-level output of the PD indicator using the Tree of Life for the birds of Kenya, considering only the persistence of species present in Kenya as relevant to maintaining Kenya’s branches of the tree, and thus quantifying their national contributions to people. As the EDGE prioritisation approach must be calculated from the global set of species to retain comparability (Isaac et al. 2007; Gumbs et al. 2018), the national-level output of the EDGE Index draws the set of EDGE species for Kenya from this global pool of priority species before recalculating all components of the EDGE Index (Figure 4b; see Appendix S2). Both indicators retain global relevance when calculated at a national level, due to the global values associated with the branches of the Tree of Life held by the species occurring in respective territories.

**Figure 4:**
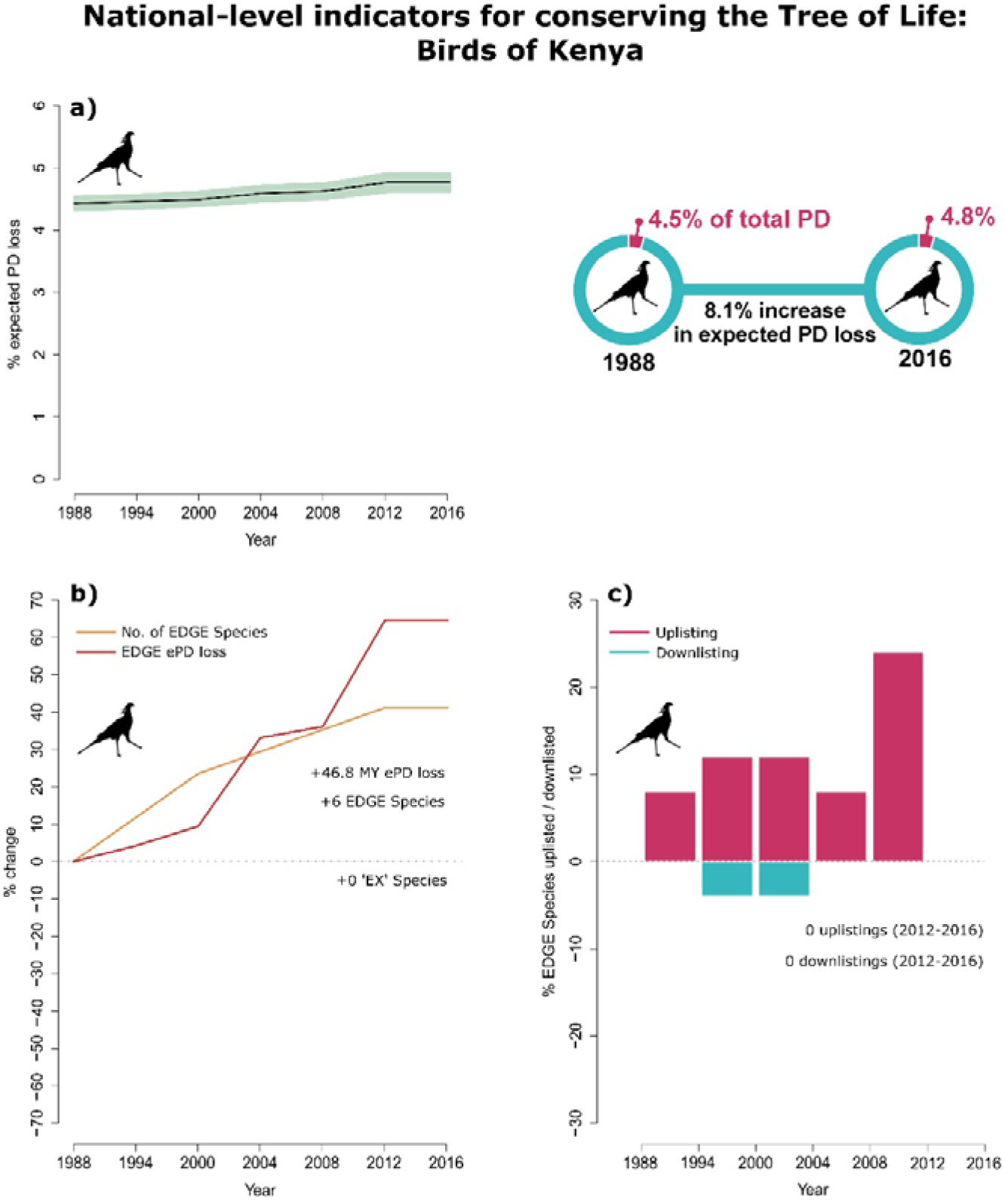
Example of national disaggregations for the two indicators for the birds of Kenya. The expected PD loss of Kenyan bird species (a) is calculated as a percentage of the total PD associated with bird species present in Kenya. The EDGE Index for Kenyan birds (b-c) is subset from the global pool of priority EDGE birds to ensure national priority species align with those of global value. See Appendix S2 for detailed methods underpinning this national disaggregation approach.

### Data availability and reporting

It is important for any indicators to be easily calculable or readily available for use by all stakeholders. For the PD Indicator, the necessary data—the evolutionary relationships of a set of species and estimates of their extinction risk—are increasingly available for large numbers of species (Hinchliff et al. 2015; Nic Lughadha et al. 2020; Cox et al. 2022). Data underpinning the initial calculation of the PD indicator for certain animal taxa by IPBES was made available as part of the commitment by the Zoological Society of London’s (ZSL) EDGE of Existence programme. The programme has been the central repository for ED and EDGE data for more than a decade) and provides regularly updated ED scores for taxonomic groups for which data are available (Gumbs et al. 2018; IPBES 2019). The Royal Botanic Gardens, Kew (RBG Kew) and partners have also committed to produce EDGE scores for the world’s seed plants (Forest et al. 2018; Royal Botanic Gardens, Kew 2021). As the EDGE of Existence programme and RBG Kew transition to generating EDGE species priorities based on principles applied here, the data underpinning the updated PD indicator and EDGE Index will be maintained and regularly updated for all animal and plant groups for which such calculations are possible.

Despite continued advances in our capability to map extinction risk across the Tree of Life (ter Steege et al. 2015; Jin & Qian 2019), more resources are needed to ensure any global biodiversity indicators are applicable to more than a narrow set of well-studied species groups, and regularly compiled to allow an effective monitoring of the current state of biodiversity. Given advances in our understanding of the Tree of Life and extinction risk for both animals (Hinchliff et al. 2015; Colston et al. 2020; Wong & Rosindell 2020; Andermann et al. 2020) and plants (ter Steege et al. 2015; Yessoufou et al. 2017; Jin & Qian 2019; Nic Lughadha et al. 2020; Borsch et al. 2020), the taxonomic scope of both indicators is set to increase between now and 2030. Beyond the baselines provided here for birds, mammals and cycads, baselines for the two indicators are also in the process of being generated for amphibians, reptiles, ray-finned fish, sharks and rays, crayfish, and various plant groups. As it becomes increasingly possible to predict extinction risk for large numbers of species (Zizka et al. 2022), the taxonomic coverage of the two indicators will remain at a minimum comparable to that of other related biodiversity indicators such as the Red List Index.

To aid the dissemination of these data and easy reporting of the proposed indicators, the IUCN Species Survival Commission’s Phylogenetic Diversity Task Force (IUCN SSC PDTF) has committed to generate both indicators at the global and national level and make these publicly available and accessible through an online tool currently in development.

## Conclusions

Conservation discourses focus on ecosystems, species and populations, but there has been an increasing recognition of the need to consider other components of biodiversity in research and policy (Pollock et al. 2017; Chaudhary et al. 2018; Mair et al. 2021), specifically variety across the Tree of Life and its importance for Nature’s Contributions to People (NCPs) (Faith 2021), as has been recognised by IPBES (IPBES 2019). The separation of Goals A (on genes, species and ecosystems) and B (on nature’s contributions to people) in the draft GBF risks isolating the conservation of biodiversity from recognition of the benefits and option values that it provides, simply assuming these values will be maintained under current conservation paradigms.

Here we have highlighted two indicators, the PD Indicator and EDGE Index, that make it practical and straightforward to incorporate the Tree of Life—and the benefits it provides— into biodiversity policy at global, regional and national scales. The indicators build on existing—and widely accepted—approaches to measuring biodiversity and extinction risk to generate monitoring tools that complement existing species-based indicators and conservation measures. The PD indicator, adopted by IPBES (Díaz et al. 2019) and updated here, has the capacity to link the preservation of biodiversity to the maintenance of benefits, bolstering intergenerational equity. The EDGE Index and its components utilise the well-established EDGE and RLI approaches to monitor the conservation status of the world’s most evolutionarily distinct species.

To be truly ambitious, global biodiversity policy must aim to safeguard the Tree of Life (Díaz et al. 2020). If we fail to do so, we risk great losses of evolutionary history including the loss of their associated options and benefits for current and future generations. But if we succeed, we can preserve much of the global value of nature’s contributions to humanity now and into the future.

