## Appendix S2 for "Conserving evolutionary history to safeguard our future: incorporating the Tree of Life into biodiversity policy"

### Supplementary Information

#### Supplementary methods

##### Phylogenetic and extinction risk data

To calculate the PD indicator and EDGE Index for mammals, we generated a set of 100 phylogenetic trees comprising all 6,253 described extant mammal species (as recognised in the Mammal Diversity Database, Burgin et al. 2018). To do this, we extracted a random sample of 100 phylogenetic trees from the distribution of mammal phylogenetic trees of Upham et al. (2019), each comprising 5,911 extant and extinct species, and imputed missing species into their respective genus at random using the ‘*congeneric.impute*’ function from the package *pez* in R (Pearse et al. 2015). We also used this approach to generate 100 phylogenetic trees for 10,988 recognised extant bird species as of January 2020 (BirdLife International 2019), imputing missing species onto a random sample of 100 trees from Jetz et al. (2014). For cycads, we subset the 100 species-level gymnosperm phylogenetic trees from Forest et al. (2018) to comprise 333 extant cycad species.

For the calculation of both the expected PD loss indicator and EDGE Index, we used IUCN Red List data to measure extinction risk. For historical extinction risk data, we used the data underpinning the Red List Index (RLI), which exists for two historical time points for cycads (2003 and 2014), seven for birds (1988, 1994, 2000, 2004, 2008, 2012, 2016), and two for mammals (1996 and 2008) and includes only genuine changes in Red List status (Henriques et al. 2020). For the PD indicator, we also included the latest available Red List data for birds (Birdlife International 2020) and mammals (IUCN 2020) to provide a provisional ‘2020’ datapoint for both groups. Following standard practice with the RLI indicator, we note that all time points for both groups are subject to change when forthcoming comprehensive RLI assessments corrects any previous assessments for all species based on new information and changes in taxonomy (Butchart et al. 2007). Both birds and mammals have had a large number of re-assessments since their most recent respective RLI time points, unlike cycads, which lack any re-assessments on the IUCN Red Lists since their most recent RLI time point in 2014 (IUCN 2020). The RLI treats species assessed as Critically Endangered (Possibly Extinct) (CR(PE)), Critically Endangered (Possibly Extinct in the Wild) (CR(PEW)), and EW as Extinct (EX), and we considered them as such for our calculations. For species lacking historical or current Red List assessments due to taxonomic discoveries/revisions, and those listed as Data Deficient (DD), we assigned them to Red List categories based on the observed distribution of Red List categories within their respective clades and repeated this for each analysis across the distributions of phylogenetic trees.

##### PD indicator

We calculated expected PD loss for each RLI time point across the 100 phylogenetic trees for birds, cycads and mammals. To calculate expected PD loss, we follow (Faith 2008; Faith et al. 2018):

$$expected PD loss=\sum_{i} \left\{ L_{i}\times\left[ \prod_{j=1}^{n_{i}} q_{j} \right] \right\}$$

Where  *i* is in index identifying individual branches, *L_i_*, is the length for branch *i, n_i_* is the number of descendant species from branch i, and q_ij_ is the extinction risk for descendant species *j* from branch *i*. To compute this in practice, we chose to multiply the length of each phylogenetic branch by the product of the probabilities of extinction of all species descendant to the branch and then calculate total PD of the resulting tree (Faith 2008). The length of the transformed branch now represents how much PD is expected to be lost in a probabilistic framework, given the probabilities of extinction of all species responsible for the persistence of the branch (the species that descend from that branch). By summing the new lengths of the branches of the resulting extinction risk-transformed tree, we can estimate the amount of PD expected to be lost across the clade based on the distribution of extinction probabilities across the species at the tips of the tree.

In lieu of accurate probabilities of extinction associated with Red List categories (Collen et al. 2016), we adopted relative weightings of extinction set to the 50-year time horizon of Mooers et al. (2008), where CR = 0.97. The RLI weightings of extinction risk, also used in the EDGE metric, represent a halving of extinction risk with each decrease in Red List category severity (Isaac et al. 2007), a pessimistic interpretation of extinction risk, which may decrease by as much as an order of magnitude for every improvement in Red List category (Butchart et al. 2004). We follow these principles and adopt the EDGE/RLI ‘halving’ approach (Isaac et al. 2007), where CR = 0.97, EN = 0.485, VU = 0.2425, Near Threatened (NT) = 0.12125, and Least Concern (LC) = 0.060625. Extinct in the Wild (EW) and Critically Endangered (Possibly Extinct; CR(PE)) were assigned a weighting of 1.

We also recalculated the expected PD loss at each time point using a pessimistic and optimistic set of extinction risk values from Mooers et al. (2008); our pessimistic extinction risk values correspond to the ‘IUCN 500’ (i.e. probability of extinction in 500 years) transformation (LC = 0.0005, NT = 0.02, VU = 0.39, EN = 0.996, CR = 1) and our optimistic extinction risk values correspond to the ‘IUCN 50’ (i.e. probability of extinction in 50 years) transformation (LC = 00005, NT = 0.004, VU = 0.05, EN = 0.42, CR = 0.97). We calculated the expected PD loss using all extinction risk values across the 100 trees for each clade (Figure S1).

Whilst the Red List Index treats all species as equal in their contributions to overall species diversity, the contribution of species to overall phylogenetic diversity varies depending on their evolutionary history. Species on long branches of the Tree of Life with few (or no) close relatives contribute more to overall PD than species on short branches with many close relatives, even more so if those relatives are threatened with extinction. By combining extinction risk data from the RLI with phylogenetic information, it is possible to also generate forecasts of best- and worst-case outcomes—and the associated expected loss—under observed and hypothetical extinction risk scenarios (Figure S2).

The PD indicator can incorporate upper and lower boundaries of expected PD loss, based on observed levels of extinction risk across a clade, to measure poor and good conservation of PD. These boundaries can provide a background against which we can measure our performance at averting the greatest losses of PD. To illustrate this, we recalculated expected PD loss for mammals at both the 1996 and 2008 time points under three different scenarios, all with the same frequency of species in each Red List category as observed in the RLI data:

1) High expected loss: Red List categories redistributed amongst species as such that the species with the highest Evolutionary Distinctiveness (ED) receive the most sever Red List categories. As ED decreases, Red List categories are assigned in descending order, from Extinct in the Wild / CR (possibly extinct) to Least Concern. Thus, higher levels of extinction risk were assigned to species with higher ED scores, and therefore greater potential contributions to expected PD loss;
2) Low expected loss: the opposite approach to (1). As the ED of species decreases, Red List categories are assigned in ascending order, from Least Concern to Extinct in the Wild / CR (possibly extinct). Thus, higher levels of extinction risk were assigned to species with the lowest ED scores, and therefore small potential contributions to expected PD loss, and the loss in the highest ED species is minimised by being assigned to LC;
3) Random expected loss: Red List categories are redistributed across mammal species at random.

We repeated this process across the 100 mammal trees to generate a distribution of scores for each scenario.

Finally, we also calculated the PD indicator using the initial IPBES methodology—for all RLI time points for cycads, birds and mammals—for comparison with our proposed updated PD indicator approach (Figure S1). Initial estimations of the IPBES PD indicator were calculated using an approximation approach that relied on published Evolutionary Distinctiveness (ED)—where branches of the Tree of Life are split equally amongst all species that descend from them (Isaac et al. 2007)—scores for sets of species (Faith et al. 2018). The approximation used the summed ED scores of threatened species (VU, EN, CR) in a clade divided by the summed ED scores of all species in the clade (equal to total PD of clade) to estimate the proportion of imperilled ED and thus approximate the expected loss of PD across the clade (Faith et al. 2018).

However, IPBES’s ED approximation does not currently take into account the interplay of shared evolutionary history and extinction risk of related species, which is important to directly calculate expected loss from branches of the Tree of Life that are shared by multiple species (Steel et al. 2007; Faith 2008). For example, an ancestral branch of the Tree of Life with two descendant species, one threatened and one non-threatened, is at less risk of being lost compared with a branch leading to two threatened species (‘PD complementarity’; Faith et al. 2004; Faith 2008). However, under IPBES’s ED approximation, each of these ancestral branches would be considered equally at risk of extinction.

Under the scenario of high expected loss (scenario 1), above), where the most severe extinction risk categories observed for mammals are redistributed to the most evolutionarily distinct species (supplementary methods), PD indicator values under our updated approach would still be lower than those reported for mammals in the IPBES Global Assessment (Figure S1, Figure S2). Further, the values from the IPBES approach are 2.2x higher for birds and 1.8x higher for mammals than previously reported estimates when the same approach to extinction risk as IPBES was taken for calculating imperilled PD within the expected PD loss framework (Gumbs et al. 2020).

##### EDGE Index

To calculate the EDGE Index we calculated the following components, for each RLI time point:

**Number of EDGE species:** Species must be both highly evolutionarily distinctive and threatened with extinction to be considered an EDGE species, thus monitoring the trends in the number of EDGE species is tantamount to monitoring the rate at which highly distinctive species are moving from non-threatened to threatened categories. A decrease in the number of EDGE species indicates that there is a reduction in the number of distinctive species facing extinction. We track trends in number of EDGE species by simply summing the number of qualifying species for each RLI time point and calculating the % increase in number of EDGE species from the earliest RLI time point available (e.g. moving from 10 to 11 EDGE Species represents a 10% increase in EDGE Species). A positive trend indicates increasing threat to the most distinctive species, and a positive % value indicates more distinctive species are in threatened categories than there were in the past. A negative % value indicates there are fewer distinctive species in threatened categories than there were in the past.

**Total expected PD loss contributions of EDGE Species:** Summing the expected PD loss contributions of all EDGE Species, or their EDGE scores, measures the amount of PD at risk of being lost across all EDGE Species, which represent some of the largest potential losses of PD. This value can increase or decrease between time points even if the number of EDGE Species remains constant: if individual EDGE Species are moving between threatened categories their expected PD loss contributions will increase or decrease as their extinction risk increases or decreases. The change in status of highly distinctive species results in a greater change in expected PD loss than a less distinct species. Therefore, these two metrics are particularly informative when used in combination. We track the trends in the expected PD loss contributions of EDGE Species as the % change from the earliest RLI time point (i.e. increasing from 100 MY to 120 MY represents a 20% increase).

**Change in extinction risk of EDGE Species:** Tracking trends in the changes in extinction risk for the most distinctive and threatened species highlights whether these species are sliding toward or moving away from extinction. This is achieved by summing the uplistings (increase in extinction risk) and downlistings (decrease in extinction risk) across all EDGE Species for each time point, and calculating the % of EDGE Species that are uplisted and downlisted between each time point. The greater the positive % value, the more species were uplisted, and the greater the negative % value, the greater the number of species that were downlisted. We track uplistings and downlistings independently to ensure they do not confound one another, and the objective would be an increasing rate of downlistings with a constant rate of zero uplistings of EDGE Species.

**Number of ‘extinct’ EDGE Species:** Hypothetically, an EDGE Species becoming extinct will lead to a reduction in the number of EDGE Species, which can appear to be a positive change in the status of EDGE Species. Therefore, monitoring the number of EDGE Species that move to one of the ‘EX’ categories (CR(PE), CR(PEW), EW, EX) allows these events to be tracked and negate any possible perversion of the other indices. To monitor this, we simply sum the number of species with above median contributions to PD (highly distinctive species) that are in one of the ‘EX’ categories at each time point. Ideally, this trend will remain constant, and may even decline with the rediscovery of species thought to be extinct.

#### Not all extinctions are equal

##### Mammal extinction scenarios and Phylogenetic Diversity loss

For all extinction-based analyses of Phylogenetic Diversity (PD) loss, we augmented a set of the 100 mammal phylogenetic trees generated for the PD indicator and EDGE Index calculations with all recently extinct mammal species (since 1500, 84 spp.; IUCN 2020). We again imputed missing species into their respective genus at random using the ‘*congeneric.impute*’ function from the package *pez* in R (Pearse et al. 2015), resulting in 100 phylogenetic trees comprising all 6,337 described extant mammal species (as recognised in the Mammal Diversity Database, 6,253 spp.; Burgin et al. 2018) and recently extinct mammal species (since 1500, 84 spp.; IUCN 2020).

To estimate the loss of PD associated with a set of mammal extinction scenarios, we measured the PD lost under each scenario as the total length of all phylogenetic branches lost when all species considered as extinct for a given scenario were pruned from the phylogeny. To estimate the mammal PD already lost to recent extinctions (our ‘past extinctions’ scenario), we calculated the amount of PD lost when the 84 species recognised as becoming extinct (EX) since 1500 were pruned from the phylogenies. For our ‘current extinctions’ scenario, we follow Rounsevell et al. (2020) in considering all mammal species recognised as Critically Endangered (Possibly Extinct) (CR(PE)) by the IUCN Red List (IUCN 2020), along with species listed as Extinct in the Wild (EW) and Critically Endangered (Possibly Extinct in the Wild) (CR(PEW)), to represent current extinction events in the process of occurring, and estimated the additional PD loss when these 31 species were pruned from each tree, following the pruning of all EX species. For the ‘future extinctions’, we calculated the additional loss in PD, following the pruning of all EX, CR(PE) and EW species, for three scenarios: (1) all 197 Critically Endangered (CR) mammal species become extinct; (2) all 701 CR and Endangered (EN) species become extinct; and (3) all 1,244 threatened mammal species—that is, CR, EN, and VU—become extinct (“No conservation” strategy; Table S1).

We then explored how averting the loss of 25% of species—a proportion we selected arbitrarily–facing extinction in each of the current and future extinction scenarios impacted the magnitude of mammalian PD loss under four simple conservation strategies: (1) 25% of threatened species selected at random (‘random’ strategy); (2) the 25% of threatened species whose conservation maximises the conservation of PD (‘PD maximisation’ strategy), identified using a greedy algorithm to maximise conservation of branches of the phylogeny (Nee & May 1997); (3) the 25% of threatened species with the smallest contributions to expected PD loss (‘low EDGE’ strategy; see EDGE Index section below for calculation of EDGE scores); (4) the 25% of threatened species with the largest contributions to expected PD loss (‘high EDGE’ strategy). We recalculated the PD loss for each extinction scenario across the 100 phylogenetic trees under the four conservation strategies to determine the amount of PD loss that is averted when equal numbers of species are conserved under different approaches (Table S1). We then calculated the percent additional PD conserved, relative to the random strategy, by the two strategies that conserved more than random (high EDGE and PD maximisation; Table S2).

***Table S1:*** *the proportion (%) of the mammalian Tree of Life lost under five conservation strategies for three future extinction scenarios (see supplementary methods). Median value is bold with range in parentheses.*

|  | Mammal phylogenetic diversity lost under each conservation strategy (%) - **median** (range) | | | | |
| --- | --- | --- | --- | --- | --- |
| **Extinction Scenario** | No conservation | Low EDGE | Random | High EDGE | PD maximisation |
| **CR** | **2.15** (1.89-2.54) | **2.02** (1.77-2.35) | **1.74** (1.48-2.03) | **1.28** (0.99-1.65) | **1.11** (0.87-1.40) |
| **CR-EN** | **6.37** (5.96-7.25) | **5.7** (5.37-6.22) | **4.39** (3.96-5.14) | **3.05** (2.63-4.2) | **2.49** (2.14-2.84) |
| **CR-VU** | **11.94** (11.42-13.07) | **10.04** (9.63-10.77) | **6.90** (6.27-7.49) | **5.53** (5.10-7.14) | **4.16** (3.78-4.62) |

***Table S2:*** *the percent increase in mammal phylogenetic diversity conserved under the high EDGE and PD maximisation conservation strategies, relative to a random conservation strategy, when the number of species extinctions remain constant. 100% = double the amount of PD conserved relative to the random conservation strategy (i.e. 100% more PD conserved; see Supplementary Methods).*

|  | Percent increase in mammalian phylogenetic diversity conserved relative to "random" strategy (%) | |
| --- | --- | --- |
| **Extinction scenario** | **High EDGE** | **PD maximisation** |
| **CR** | 109.99 | 151.64 |
| **CR-EN** | 67.78 | 96.43 |
| **CR-VU** | 27.19 | 54.30 |

##### National disaggregation

It is important for any biodiversity indicators incorporated into the GBF to be applicable at regional or national scales(Secretariat of the Convention on Biological Diversity 2021b). To demonstrate the capability of the two indicators we present here in meeting the needs of nations to monitor their progression towards the framework’s Goals and Targets, we present a national disaggregation for each of them. We used birds for this example as they are the only taxonomic group with more than two Red List Index time points available, thus provide a more detailed illustration of changes through time. We selected Kenya due to its high richness in both bird species and EDGE species (24 EDGE bird species). A flow diagram indicating the main steps of each calculation is presented in Figure S3.

**Phylogenetic Diversity indicator:** There are several possible approaches for calculating the national disaggregation of the PD Indicator, two of which we consider as feasible now for reporting. To generate the national-level expected PD loss indicator for Kenyan birds presented here, we simply pruned the 100 bird phylogenetic trees generated for the global calculation, with Kenyan bird species determined using presence data provided by the IUCN Red List API in R (IUCN 2020) (Figure S 2). In other words, for each of the 100 phylogenetic trees produced with all species at a global scale, we kept only the species occurring in Kenya. We then applied the same method as with the global-level indicator to calculate the percent of total PD associated with Kenyan species (i.e. summed branch lengths of the pruned trees) expected to be lost given to the extinction risk of Kenyan species.

This approach does not consider the influence of the extinction risk of species that are not present in Kenya but which descend from internal branches shared with species present in Kenya. For example, under this approach the expected loss of an internal branch with two descendant species, one in Kenya, one in Tanzania, is not influenced by the persistence of the species in Tanzania, which may be of low extinction risk. This approach assumes that each individual country takes responsibility for the maintenance of internal branches within its borders.

The alternative approach, where the extinction risk of the descendant species found only in Tanzania influences the persistence of the internal branch in Kenya, naturally reduces Kenya’s responsibility to maintain the internal branch within its borders as it persists elsewhere. This alternative approach presents the problems of reducing national responsibility for the maintenance of biodiversity and of providing a misleading positive result for nations that do not improve the extinction risk of any species but share internal branches with other nations, as they will appear to be improving the national status of their biodiversity and performing well.

We here present the first approach, which provides a measure of the biodiversity present within national borders representing that country’s nature’s contributions to people. In this scenario it is the responsibility of the nation in question to maintain that biodiversity to maximise nature’s contributions to people both within their borders and at a global scale. However, this national-level calculation can not be directly derived from the global calculation of the indicator, which may be a more desirable attribute. The second approach, which factors in the influence of descendant species of branches found in a country that are not present in that country, can be easily disaggregated from the global PD indicator dataset without recalculations and may indeed be more desirable for monitoring the status of the Tree of Life at the national level. Both approaches have associated benefits and limitations, and thus we commit to producing one or both as needed for policy reporting.

**EDGE Index:** The EDGE scores of species are global in the sense that they are calculated from the complete Tree of Life for the focal taxonomic group (Isaac et al. 2007; Gumbs et al. 2018). EDGE scores, and thus relative rankings of species within a taxonomic group, are therefore calculated at the global level before being disaggregated to subsets of species present in a given country (Figure S2). This means that species that co-occur in multiple countries retain consistent relative EDGE rankings, and all national-level priority EDGE species must also therefore be priority species at the global level. To calculate the national-level EDGE Index for Kenyan birds, we subset the global EDGE Index for birds to just those species present in Kenya. We then recalculated each element of the Index (extinctions, change in expected PD loss and number of EDGE species, and changes in Red List category through time) as a function of the total number of EDGE species present in Kenya, rather than the global EDGE species pool, as with the global EDGE Index.

Though not applied here, it is possible to use the national—in favour of global—RLI approach (Rodrigues et al. 2014) to monitor trends in the extinction risk of both EDGE species and PD. Recent work highlights the scale-dependent nature of biodiversity patterns and endemism, particularly with regard to distributing branches of the Tree of Life in space (Jarzyna & Jetz 2018; Daru et al. 2020), and this scale dependency needs to be considered in any application of the national RLI approach. As EDGE rankings are inherently global due to their calculation from an entire clade (Gumbs et al. 2018), and the aim of the EDGE Index is not to generate a single trendline (as is the case with national RLIs), the application of the national RLI approach is not directly applicable to national EDGE Index calculations.

##### Identifying evolutionarily distinct species

In 2012, IUCN adopted a resolution that recognised the importance of conserving threatened evolutionarily distinct lineages and their irreplaceable traits and genes (IUCN 2012). Díaz et al. (2020) echoed IUCN’s concerns when they recommended that any ambitious post-2020 targets must include an element to ensure the conservation of evolutionarily distinct species in order to maintain the full breadth of evolutionary history. To achieve this, Díaz et al. (2020) proposed a novel approach to identify such evolutionarily distinct species (annotation (h), supplementary materials) whereby species that do not have multiple close relatives are prioritised. In their paper, these species are defined as those with more than one related species that descended from the same common ancestor within the timespan of the average species age +/- 1 MY.

There are, however, several existing approaches for measuring the evolutionary distinctiveness of species, three of which are widely established in global conservation research: 1) the ‘fair proportion’ (‘FP’) metric (Redding & Mooers 2006), which provided the evolutionary distinctiveness measure for the original EDGE metric (Isaac et al. 2007); 2) the expected PD contribution of a species, or it’s Heightened Evolutionary Distinctiveness (‘(H)ED’; Steel et al. 2007), which underpins the latest iteration of the EDGE metric utilised by the Zoological of Society of London’s EDGE of Existence programme (Gumbs et al. 2022); and 3) the terminal branch length (‘TBL’) of a species, which is its minimum unique contribution to the Tree of Life (Faith et al. 2004; Gumbs et al. 2020).

To test the performance of the proposed approach of Díaz et al. (2020) at maintaining the Tree of Life, we calculated the set of species matching the Díaz et al. (2020) criteria for evolutionarily distinct species across our 100 mammal phylogenetic trees. For each tree we then calculated FP, (H)ED and TBL metrics for all species and selected the highest-ranking sets of priority species for each that were equal in number to the species identified under Díaz et al.’s approach. We then calculated, for each tree, the percent of total PD (i.e. proportion of Tree of Life; Figure S3a) and threatened PD (i.e. PD to be lost if all VU, EN, CR species become extinct; Figure S3b) captured by priority species under each metric. Díaz et al.’s proposed measure of evolutionary distinctiveness captured significantly less total PD compared with all established methods (p < 0.0001 following Tukey’s honest significant differences); captured significantly less threatened mammalian PD than (H)ED and TBL (p < 0.0001); and captured comparable amounts of threatened mammalian PD to the FP metric (p = 0.94).

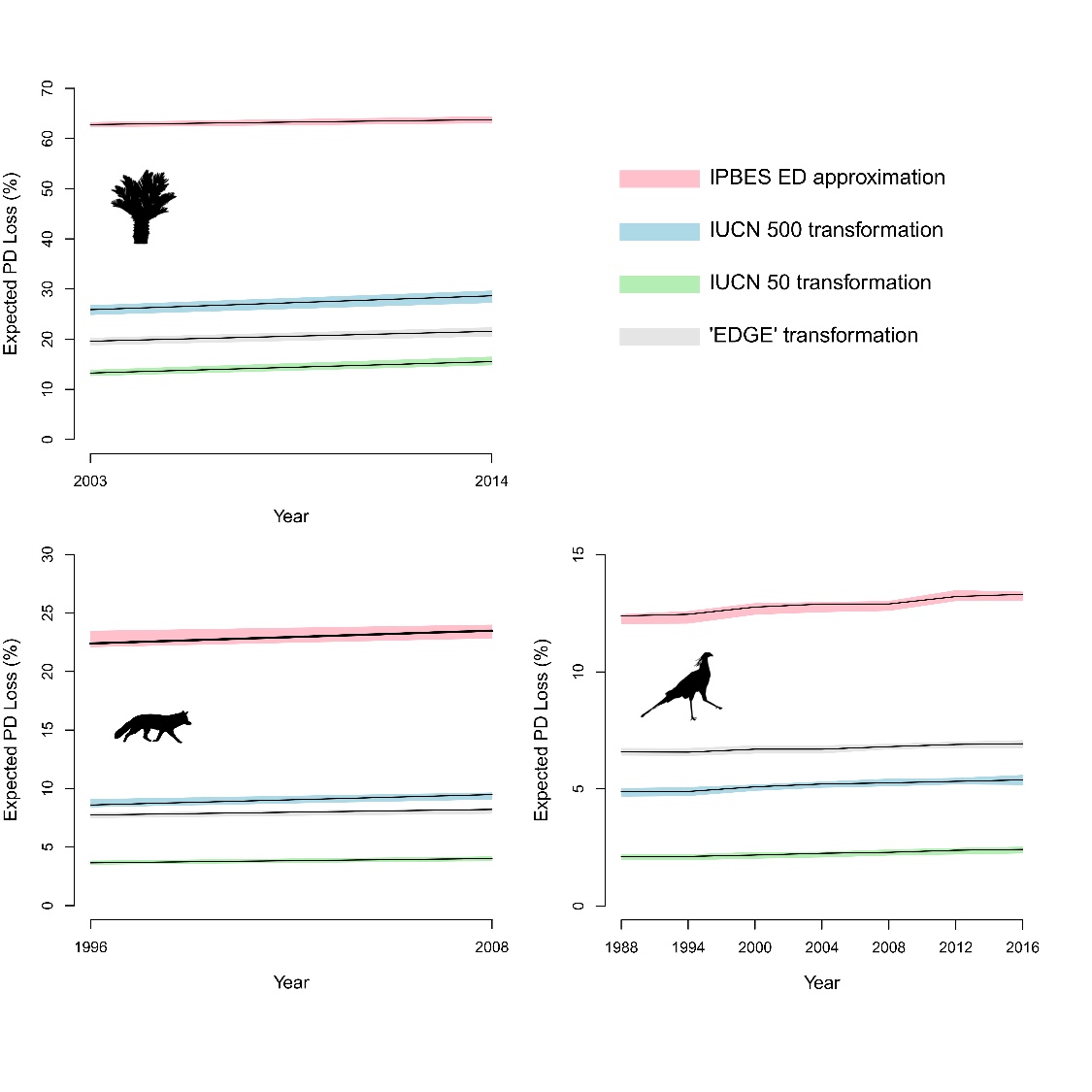

***Figure S1****: The PD indicator under different extinction risk quantifications. The expected PD loss for cycads (top left), mammals (bottom left) and birds (bottom right) at published RLI timepoints under four quantifications of extinction risk. The shaded regions around each trend line represent the range of values.*

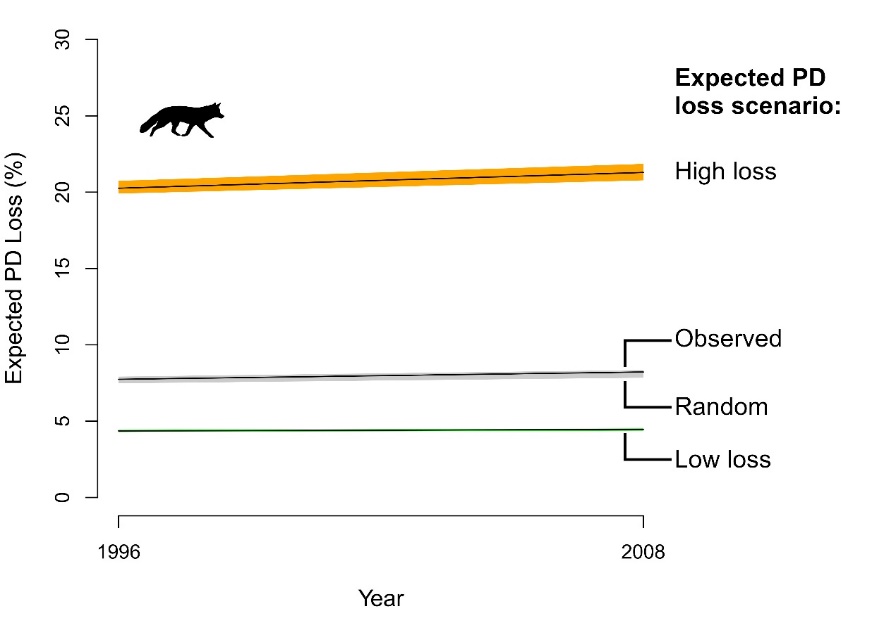

***Figure S2:*** *Expected PD loss scenarios. Variation in expected PD loss for mammal for RLI timepoints, under four scenarios of species loss, where the current observed frequency of Red List categories is maintained: ‘High loss’ = most severe Red List categories redistributed to species with highest ED scores; ‘Low loss’ = most severe Red List categories redistributed to species with lowest ED; ‘observed’ = Red List categories assigned as currently listed by RLI; ‘Random’ = Red List categories randomly distributed across species 100 times. All scenarios calculated across 100 phylogenetic trees.*

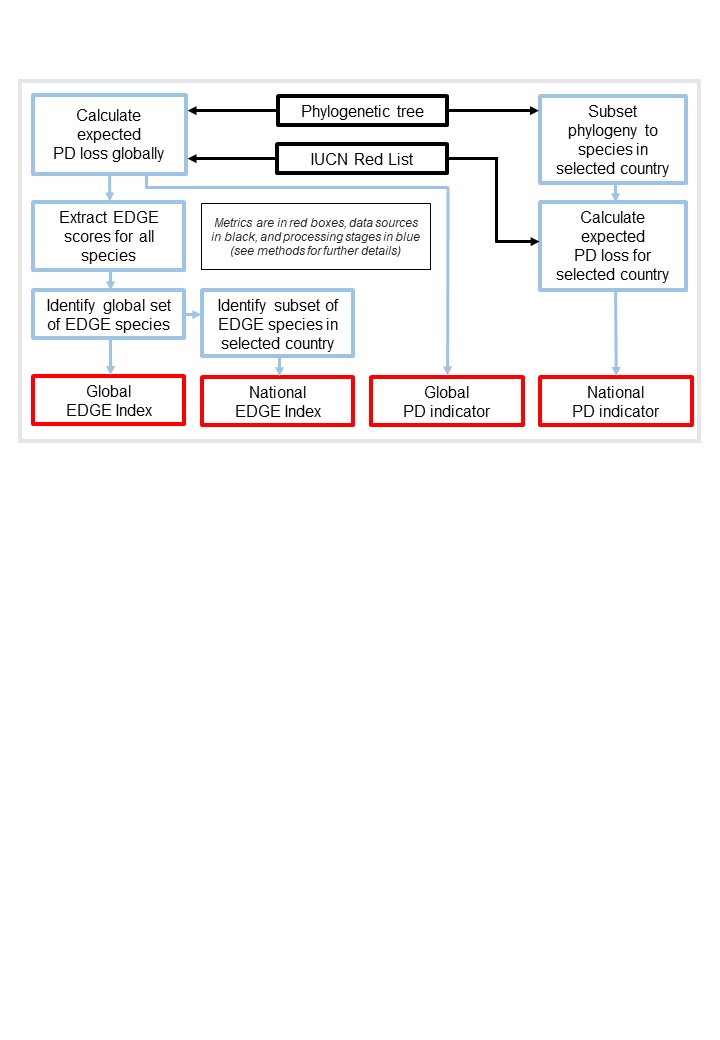

***Figure S3:*** *The process to generate global and national-level PD-based indicators. The input data (black), calculation and data processing stages (blue) and indicator outputs (red) are detailed in the text of these supplementary methods.*

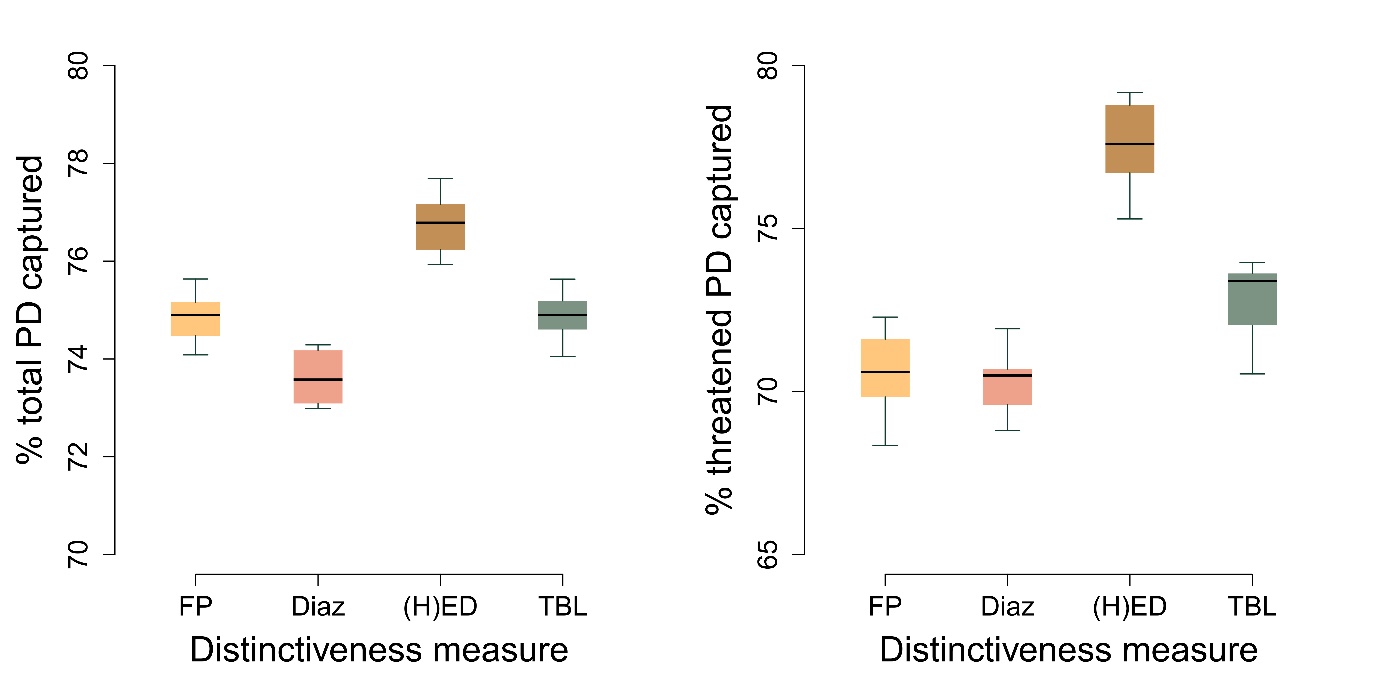

***Figure S4:*** *Performance of various evolutionary distinctiveness measures at capturing (a) total and (b) threatened mammalian Phylogenetic Diversity (PD). Species that meet D*í*az et al. (2020)’s criteria for evolutionary distinctiveness were identified for 100 mammal trees and sets of equal numbers of species were identified from the highest-ranking species for each other metric: Fair Proportion (FP), which underpins original ED calculations; Heightened Evolutionary Distinctiveness ((H)ED) which underpins the EDGE approach here; and Terminal Branch Length (TBL), which is the minimum unique contribution of each species to total PD. The percent of total and threatened PD these sets of species capture was calculated for each tree.*

***Table S3:*** *Change in expected PD loss for each clade under each extinction risk quantification.*

|  | Expected PD loss (initial exp. PD loss – latest exp. PD loss **(relative change)**) | | | |
| --- | --- | --- | --- | --- |
| Clade | IPBES binary | IUCN 50 | IUCN 500 | EDGE |
| Birds | 12% - 13% **(7.5%)** | 2.1% – 2.4% **(15%)** | 4.7% - 5.2% **(10.3%)** | 6.6% – 6.9% **(5.4%)** |
| Cycads | 62.8% - 63.8% **(1.5%)** | 13.8% - 15.5% **(17.1%)** | 25.9% - 28.6% **(10.7%)** | 19.6% - 21.6% **(10%)** |
| Mammals | 22.4% - 23.5% **(4.9%)** | 3.7% - 4% **(10%)** | 8.6% - 9.5% **(10.3%)** | 7.7% - 8.2% **(6.2%)** |
