## Appendix S1 for "Conserving evolutionary history to safeguard our future: incorporating the Tree of Life into biodiversity policy"

### The tree of life: Conserving our evolutionary heritage to ensure NCPs for future generations

At the heart of the post-2020 GBF is the recognition that we must value and maintain Nature's Contributions to People, both now and in the future to ensure intergenerational equity. If global policy is to 'bend the curve' of biodiversity loss whilst securing a broad range of benefits, we must set and attain highly ambitious goals that include prioritising the conservation of evolutionarily distinct lineages to effectively safeguard the Tree of Life[1].

By conserving the Tree of Life, we can secure both the benefits and future options for humanity provided by biodiversity, capturing non-monetary benefits and ensuring intergenerational equity, as encompassed by NCPs. However, the proposed headline indicators to be used for Goal B relate only to physical monetary ecosystem services and assets, or our impact on the planet, and does not capture benefits from biodiversity at all. This neglects an entire set of non-monetary benefits and options that biodiversity provides, which must be secured. Further, current species-focused indicators capture information per species but do not account for between-species diversity and the features and traits that arise as a result of our planet's evolutionary history. We fill this gap with two indicators to monitor our progress towards safeguarding the Tree of Life, and associated benefits, that can address these omissions, and uniquely link Goals A and B.

We can effectively maintain nature's contributions to people by conserving evolutionary history – encompassing the variety of characteristics that have emerged amongst species through evolution, and which bestow nature's benefits. We can now predict that the loss of evolutionary history will have strong negative impacts on the benefits and future options for humanity provided by biodiversity.

Averting these losses requires considering the diversity across the Tree of Life —the evolutionary relationships of all living species — and ensuring that deep branches of the Tree are conserved. This means protecting species with many millions of years of unique evolutionary history, such as the Ginkgo tree or the Red Panda, as well as areas of the Tree of Life with unique lineages of species that are all threatened, such as pangolins, elephants, or rhinoceroses.

Overlooking the biodiversity-based measures of NCPs such as evolutionary history, in Goal B and the post-2020 GBF more widely, neglects the role of biodiversity in the provision of benefits and the role of conservation in maintaining those benefits for the future.

The Tree of Life is at risk across the planet, and the species whose conservation would avert the greatest losses, such as EDGE species, are found in all regions and all countries, from seabirds across Northern Europe to the distinctive parrots of New Zealand, and the endemic flora of Chile.

**Box 1**: Only the measure Phylogenetic Diversity[2], a critical but often overlooked facet of biodiversity, can account for between-species diversity and the features and traits that arise as a result of our planet's evolutionary history. This diversity is an essential basis for biodiversity's provision of benefits to people.

- **Expected loss of Phylogenetic Diversity (PD Index)[3]**

For Goal B, the Phylogenetic Diversity indicator monitors biodiversity's capacity to provide benefits into the future, and is already used by IPBES to monitor multiple NCPs, including the provision of medicinal and biochemical resources, and the maintenance of options for the future that biodiversity provides[4].

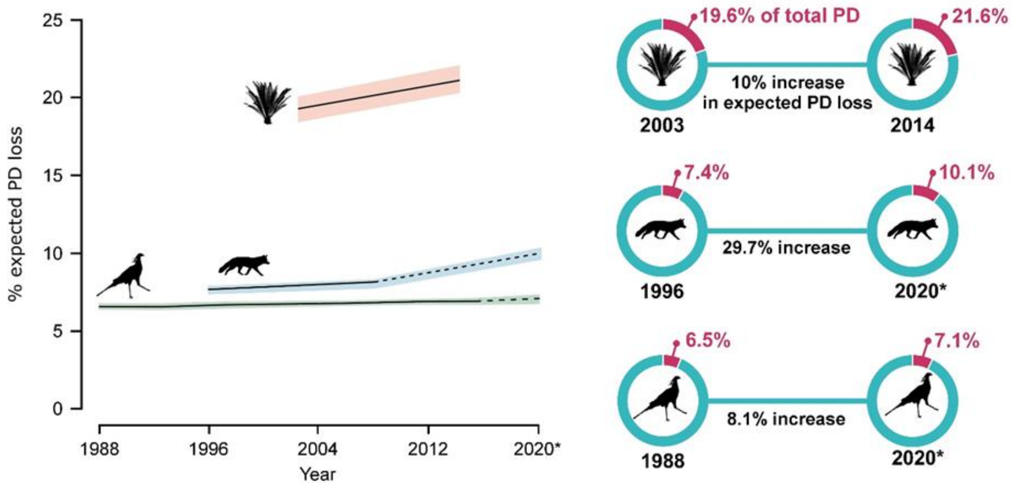


*Figure 1: The PD indicator: tracking expected PD loss through time. Left panel: trends in percentage of expected PD loss for the world's mammals (blue), birds (green) and cycads (pink), based on current and historical IUCN Red List assessments; right panel: detail of this change, baseline (left circle) and latest (right circle) estimations of expected PD loss for each clade, with the percent change in overall expected PD loss.*

[1] Diaz et al. 2020
[2] Faith 1992
[3] Gumbs et al. 2022
[4] Faith et al. 2018

**Box 2:** A practical methodology to apply PD to conservation is embodied in the EDGE lists produced by the Zoological Society of London (ZSL). EDGE (Evolutionarily Distinct and Globally Endangered) species are those which represent billions of years of threatened evolutionary history[5], major opportunities to avert loss of PD and the associated loss of options, alongside their heritage and existence values as highly distinctive species[6].

- **Changing status of Evolutionarily Distinct and Globally Endangered species (EDGE Index)[3]**

For Goal A, the component EDGE (Evolutionarily Distinct and Globally Endangered) Index monitors how well we are performing at averting the greatest losses across the Tree of Life by conserving the most distinctive species.

These two indicators create a formal linkage between goal B on benefits from biodiversity, with the conservation of species in Goal A, which is currently absent - in comparison to the existing links between area conservation and valuing ecosystem services. This omission risks prioritising conservation activities for maintaining ecosystem services while assuming that sufficient biodiversity will also be conserved.

***Data availability & Country-level disaggregations***

The IUCN SSC's Phylogenetic Diversity Task Force has committed to generate both indicators at the national and global levels (currently for jawed vertebrates, gymnosperms, and crayfish, and working to expand this across the tree of life), via open source portals to enable rapid and easy national-level reporting for Parties, requiring no additional effort or capacity needs.

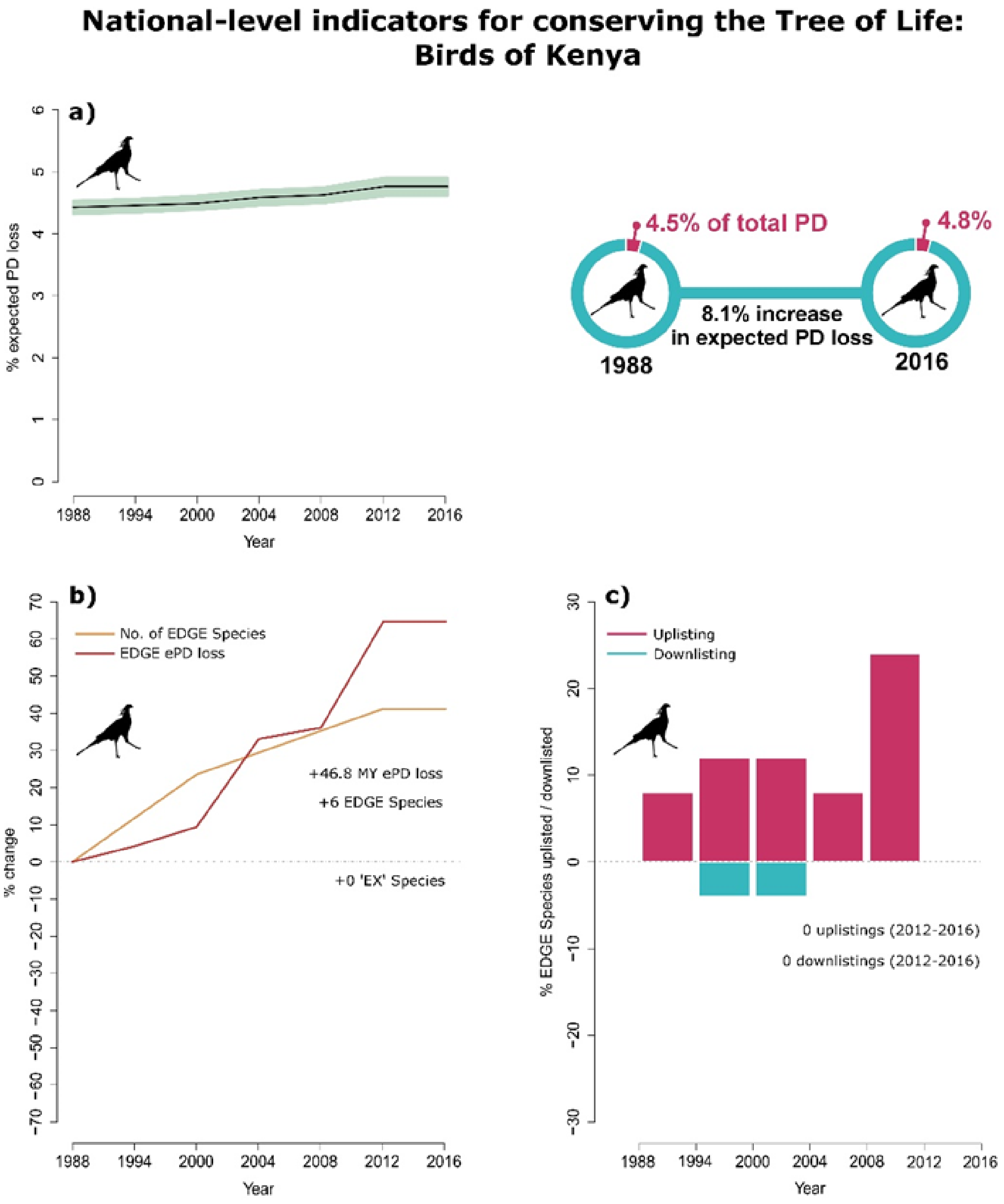


***Figure 2:*** *Example of national disaggregations for the two indicators for the birds of Kenya. The expected PD loss of Kenyan bird species (a) is calculated as a percentage of the total PD associated with bird species present in Kenya.*

*The EDGE Index for Kenyan birds (b-c) is subset from the global pool of priority EDGE birds to ensure national priority species align with those of global value. Left panels: tracking changes through time in the total number of EDGE species, associated expected PD loss (ePD loss), and extinctions (EX Species), of priority EDGE Species per clade; and (right panels) the changes in extinction risk (uplistings and downlistings: species moving into higher or lower Red List categories) within sets of EDGE Species.*

[5] Gumbs et al. 2020

[6] Owen et al. 2019

*The IUCN SSC Phylogenetic Diversity Task Force, hosted by On the EDGE Conservation, provides guidance and expertise on phylogenetically informed conservation. Find out more at www.pdtf.org and www.ontheedge.org*
*Disclaimer - The opinions given herein belong solely to the PDTF and do not represent the views or policies of IUCN*
